# Chromosomal scale length variation of germline DNA can predict individual cancer risk

**DOI:** 10.1101/303339

**Authors:** Chris Toh, James P. Brody

## Abstract

Inherited factors are thought to be responsible for a substantial fraction of many different forms of cancer. However, individual cancer risk cannot currently be well quantified by analyzing germ line DNA. Most analyses of germline DNA focus on the additive effects of single nucleotide polymorphisms (SNPs) found. Here we show that chromosomal-scale length variation of germline DNA can be used to predict whether a person will develop cancer. In two independent datasets, the Cancer Genome Atlas (TCGA) project and the UK Biobank, we could classify whether or not a patient had a certain cancer based solely on chromosomal scale length variation. In the TCGA data, we found that all 32 different types of cancer could be predicted better than chance using chromosomal scale length variation data. We found a model that could predict ovarian cancer in women with an area under the receiver operator curve, AUC=0.89. In the UK Biobank data, we could predict breast cancer in women with an AUC=0.83. This method could be used to develop genetic risk scores for other conditions known to have a substantial genetic component and complements genetic risk scores derived from SNPs.

## Introduction

Different lines of evidence suggest that a large fraction of common cancers are due to inherited factors. Cancer rates in monozygotic and dizygotic twins differ(1). Similarly, age-specific incidence studies suggest that most cancers occur in a small subset of the population, probably determined by inherited genetics (2).

Genome wide association studies (GWAS) can identify inherited genetic factors that influence phenotypes(3). GWAS have been very successful at identifying single DNA alterations that influence traits, but if a trait requires two or more independent DNA alterations, known as epistatic interactions, it is generally invisible to GWAS using the typical statistical analysis(4,5).

Machine learning can take many weak predictors and create a strong ensemble classifier, combining these predictors in non-linear fashion; these algorithms should be able to identify epistatic interactions. However, most machine-learning algorithms require more observations than predictors. Hence, one cannot effectively apply machine learning algorithms to GWAS data when the number of observations, known as cases and controls (typically less than 10,000), is less than the number of predictors, or single nucleotide polymorphisms, observed (typically half a million or more).

The best current genetic predictive tests use a polygenic risk score computed by adding the risk from many different single nucleotide polymorphisms (SNPs). In 2015, a large collaboration published a study demonstrating that a polygenic risk score based on 77 SNPs could stratify breast cancer risk in women. They found that women who scored in the top 1% had a three-fold increase in risk compared to women who scored in the middle quintile (6).More recently, a group from Myriad Genetics developed a risk score based upon 82 different SNPs and applied it to a group of women who have a family history of breast cancer but lack monogenic mutations in well-known breast cancer genes. They found a significant increase in breast cancer among those women with the highest risk scores(7). Another recent study computed genome wide polygenic scores to predict risk in five common diseases, including breast cancer, in the UK Biobank data using several million different SNPs(8).

To identify inherited disease caused by epistatic interactions with machine learning one needs to identify a genetic measurement that condenses the information contained within the genome into fewer numbers, while maintaining sufficient information about genetic differences that characterize differences among humans. Chromosomal scale length variation satisfies these criteria. It condenses multiple smaller length variants known as copy number variants, insertions, deletions or duplications, by simply adding repeats and subtracting deletions to the overall length. Here we test whether a dataset that characterizes humans by a set of chromosomal scale length variation measurements can be used to find an accurate measure of individual risk for developing particular cancers.

## Results

The Cancer Genome Atlas (TCGA) was a project sponsored by the National Cancer Institute to characterize the molecular differences in 33 different human cancers (9,10). The project collected samples from about 11,000 different patients, all of whom were being treated for one of 33 different types of tumors. The samples collected usually included tissue samples of the tumor, tissue samples of normal tissue adjacent to the tumor and normal blood samples.

We used the TCGA data on copy number variation from samples extracted from the patient’s peripheral blood (normal blood samples). This data had been processed by TCGA through a bioinformatic pipeline that began with microarray measurements and concluded with Birdsuite, a software package that identifies copy number variation. Birdsuite identified many typical copy number variations of a few thousand bases long, but also identified some very long stretches that encompass over 90% of each chromosome, which we call measurements of chromosomal-scale length variation.

For each type of tumor, we created a case/control study. Cases were the number of “normal blood” samples from patients with those diagnoses. Controls were the number of “normal blood” samples from patients without that specific diagnosis.

We set up as a binary classification supervised learning task to distinguish between patients diagnosed with one cancer (breast cancer, for example) and those not diagnosed with that cancer. Figures 1 and Table 1 present the overall results.

**Fig. 1.**
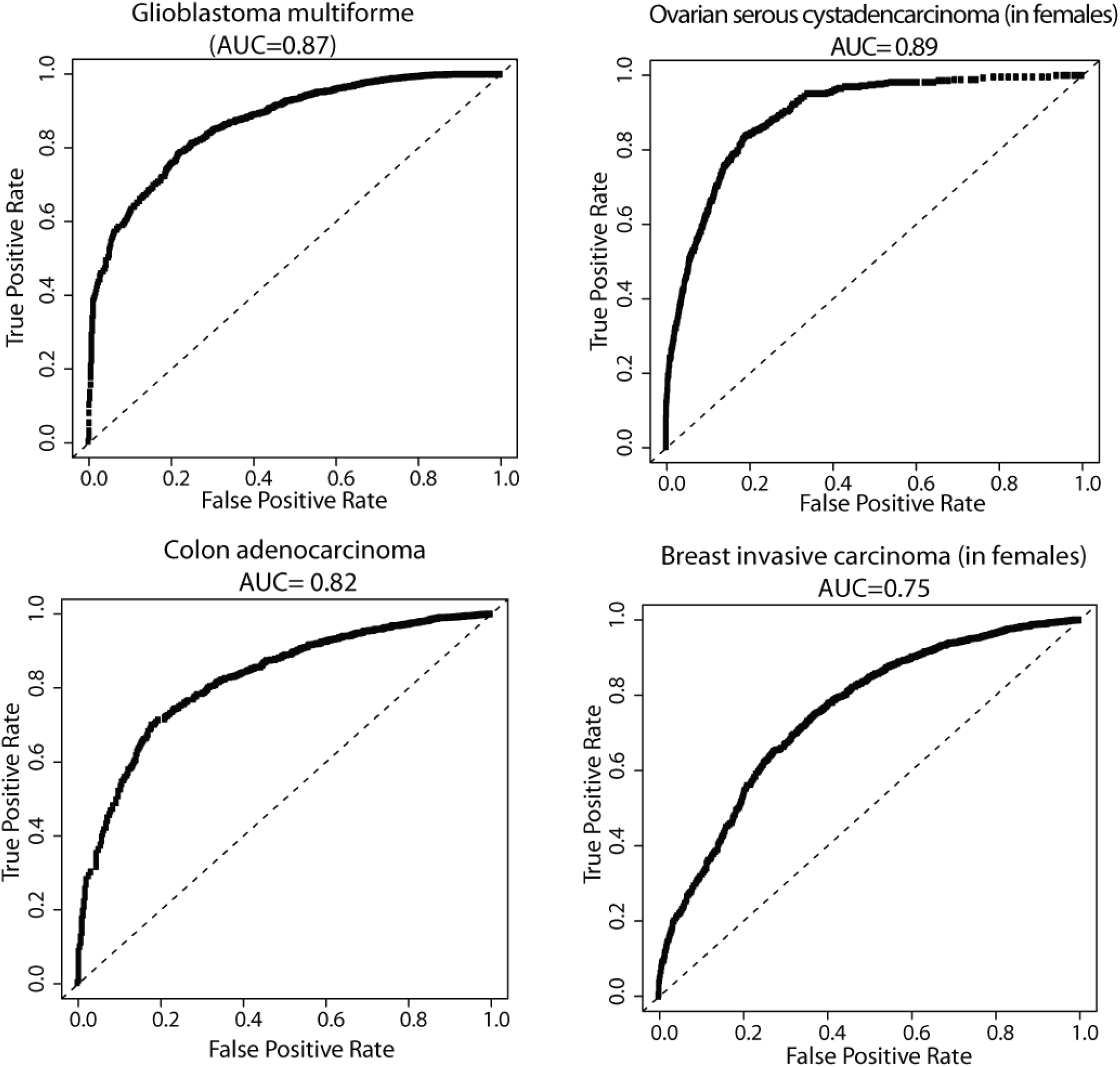
The receiver operator curves for four different cancer types. The receiver operator curves characterize how the test’s true positive rate varies with its false positive rate. The overall curve can be summarized by the area under the curve (AUC). An AUC of 0.50 is indicative of a random guess. An AUC of 1.0 is a perfect test; all cancer cases and controls are correctly classified.

**Table 1.** This table shows the cancer type, number of batches, area under the receiver operator curve (AUC), the number of cases in the TCGA dataset, and the number of controls in the TCGA dataset. The average of five measurements is reported. With one exception, the standard deviation of the five measurements was less than 0.03, usually less than 0.01. The standard deviation of kidney chromophobe’s five measurements was higher, 0.10, probably because there were only nine cases of kidney chromophobe in the dataset.

| Type of Cancer | Batches | Cases | Controls | AUC |
| --- | --- | --- | --- | --- |
| Adrenocortical carcinoma | 2 | 85 | 9713 | 0.77 |
| Bladder Urothelial Carcinoma | 23 | 361 | 9437 | 0.70 |
| Breast invasive carcinoma (in women) | 37 | 968 | 3715 | 0.73 |
| Breast invasive carcinoma (in men and women) | 37 | 977 | 8821 | 0.81 |
| Cervical squamous cell carcinoma | 19 | 281 | 4411 | 0.71 |
| Cholangiocarcinoma | 2 | 34 | 9764 | 0.71 |
| Colon adenocarcinoma | 20 | 374 | 9424 | 0.79 |
| Lymphoid/ Diffuse Large B-cell Lymphoma | 6 | 42 | 9756 | 0.76 |
| Esophageal carcinoma | 10 | 124 | 9674 | 0.78 |
| Glioblastoma multiforme | 19 | 484 | 9314 | 0.86 |
| Head and Neck squamous cell carcinoma | 18 | 488 | 9310 | 0.66 |
| Kidney Chromophobe | 3 | 9 | 9789 | 0.71 |
| Kidney renal clear cell carcinoma | 11 | 110 | 9688 | 0.82 |
| Kidney renal papillary cell carcinoma | 16 | 219 | 9579 | 0.75 |
| Brain Lower Grade Glioma | 16 | 485 | 9313 | 0.66 |
| Liver hepatocellular carcinoma | 19 | 306 | 9492 | 0.69 |
| Lung adenocarcinoma | 20 | 402 | 9396 | 0.67 |
| Lung squamous cell carcinoma | 21 | 296 | 9502 | 0.72 |
| Mesothelioma | 2 | 85 | 9713 | 0.82 |
| Ovarian serous cystadenocarcinoma | 15 | 424 | 4268 | 0.89 |
| Pancreatic adenocarcinoma | 11 | 146 | 9652 | 0.70 |
| Pheochromocytoma and Paraganglioma | 1 | 175 | 9623 | 0.84 |
| Prostate adenocarcinoma | 18 | 423 | 3711 | 0.66 |
| Rectum adenocarcinoma | 10 | 132 | 9666 | 0.75 |
| Sarcoma | 15 | 233 | 9565 | 0.72 |
| Skin Cutaneous Melanoma | 14 | 465 | 9333 | 0.73 |
| Stomach adenocarcinoma | 17 | 369 | 9429 | 0.74 |
| Testicular Germ Cell Tumors | 1 | 132 | 4002 | 0.73 |
| Thyroid carcinoma | 17 | 417 | 9381 | 0.73 |
| Thymoma | 1 | 111 | 9687 | 0.78 |
| Uterine Corpus Endometrial Carcinoma | 25 | 504 | 4188 | 0.71 |
| Uterine Carcinosarcoma | 1 | 48 | 4644 | 0.78 |
| Uveal Melanoma | 1 | 80 | 9718 | 0.80 |

Table 1 shows the cancer type, number of batches, area under the receiver operator curve (AUC), the number of cases in the TCGA dataset, and the number of controls in the dataset. The average of five measurement is reported. With one exception, the standard deviation of the five measurements was less than 0.03, usually less than 0.01. The standard deviation of kidney chromophobe’s five measurements was higher, 0.10, probably because the dataset only contains nine cases of kidney chromophobe.

Figure 1 presents the receiver operator curves for four of the tumor types. These four were selected for different reasons. Gliobastoma multiforme (11) and ovarian serous cystadenocarcinoma (12) had the largest AUC values, while colon adenocarcinoma (13) and breast invasive carcinoma (14) were chosen because they are the most common types of cancer.

Table 2 presents the results of an example test to predict glioblastoma multiforme in patients. A test dataset consisting of 20% of the TCGA data was set aside. A machine-learning model was trained using the other 80% of the data. The model was then applied to the test data. We found that we could predict about 10% of these brain cancers without any false positives.

**Table 2.**
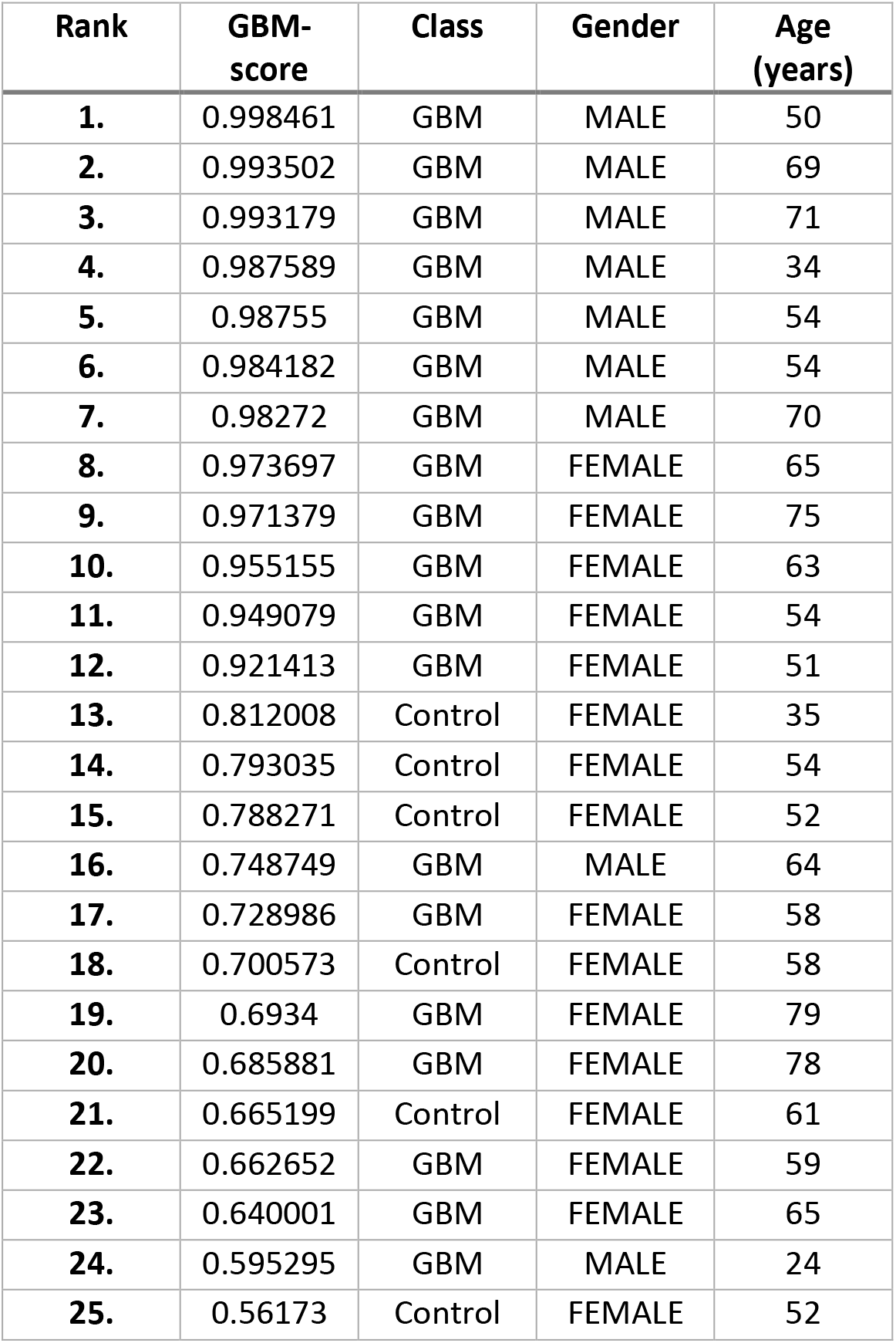
A machine-learning ranking for glioblastoma multiforme (GBM). The TCGA dataset was split into a training/validation set (80%) and a test set (20%). The model was built with the training/validation set and then applied to the test set. The test set contained information from 1704 patients. Of these, 101 patients had been diagnosed with glioblastoma multiforme and 1603 had not. The 1603 were designated as controls. The model computes a GBM-score based on the patient’s gender and chromosome-scale length variation. The patient’s age is not used in the model, but is provided here. This table lists the top 25 patients, ranked by the model’s GBM-score. The probability that the top 12 were all drawn from the GBM class by chance is about one in 10^14^.

We also tested the effect of race on these predictions. The category “race” of patients in the TCGA data is listed as “white” (6,406 patients), “black or African American” (776 patients), or “Asian” (603 patients). A very small number are listed as “American Indian or Alaska native” (20 patients) or “native Hawaiian or other Pacific Islander” (11 patients). “Unknown or Not available” was listed for 1,010 of the 8,826 patients. We constructed a training set consisting of 80% of the patients with a racial category listed as “White.” We then built models to predict ovarian cancer, breast cancer, colon cancer and glioblastoma multiforme using data from only white patients. We tested the predictive ability of these models on three different datasets of patients identified as white, black or African American, and Asian. The breakdown for these datasets is shown in Table 3. The results of these predictive tests are shown in Table 4.

**Table 3.** The racial composition of the cases and controls for four types of cancer in the TCGA dataset.

|  | Ovarian |  | Glioblastoma<br>Multiforme |  | Breast Cancer |  | Colon Cancer |  |
| --- | --- | --- | --- | --- | --- | --- | --- | --- |
|  | Case | Control | Case | Control | Case | Control | Case | Control |
| **White** | 359 | 3098 | 422 | 5984 | 652 | 2805 | 196 | 6210 |
| **Black or Afr. Am.** | 25 | 496 | 45 | 731 | 163 | 358 | 57 | 719 |
| **Asian** | 16 | 261 | 11 | 592 | 59 | 218 | 11 | 603 |

**Table 4.** We trained a model with TCGA data exclusively from patients labelled as “white.” A portion (20%) of the white patients was withheld from the training set. The model was then applied to three different validation datasets from the TCGA data. The first contained the withheld 20% of the white patients, the second contained all patients labelled Black or African American and the third contained all patients labelled Asian. The model trained on white patients consistently performed better on Asian patients than on Black/African American patients.

| Validation | Ovarian | Glioblastoma Multiforme | Breast Cancer | Colon Cancer |
| --- | --- | --- | --- | --- |
|  | AUC | AUC | AUC | AUC |
| **White** | 0.93 | 0.82 | 0.74 | 0.81 |
| **Black or Afr. Am.** | 0.70 | 0.73 | 0.56 | 0.57 |
| **Asian** | 0.98 | 0.88 | 0.83 | 0.81 |

We then replicated this process in an independent dataset, the UKBiobank dataset. The UKBiobank project consists of half a million people aged between 40-69 years recruited between the years 2006-2010(15). These people volunteered for the study and are healthier than the general population(16). Most have supplied biological samples and filled out questionnaires about their health. In addition, their medical records have been linked with national cancer registries. Germ line DNA samples were extracted from these samples and analyzed with the UK Biobank Axiom Array by the UK Biobank. The UK Biobank dataset that we used consisted of measurements at 820,967 genetic markers across 23 chromosomes for each of 488,377 different patients.

From the UK Biobank population, we identified 1,534 women who both self-reported having been diagnosed with breast cancer and were identified by cancer registries as having been diagnosed with breast cancer. We used a control population of 4,391 cancer-free women.

We quantified the genetics of each of these women with 88 numbers, each representing the length variation of one quarter of each of 22 chromosomes. We did not use the X chromosome. Using this labelled dataset, consisting of 5,925 women with 88 measurements for each, we found a classifier with an AUC of 0.83 for identifying breast cancer.

We also used the UKBiobank dataset to better understand chromosomal-scale length variation. We computed the correlations between the 88 different measurements using a dataset of 10,000 people from UKBiobank and plotted the chromosomal-scale length variation for different segments of the genome to visualize the correlations. Figure 2 demonstrates that the correlation between chromosomal-scale length variation of different genomic locations measured for many different subjects is highly variable.

**Fig. 2.**
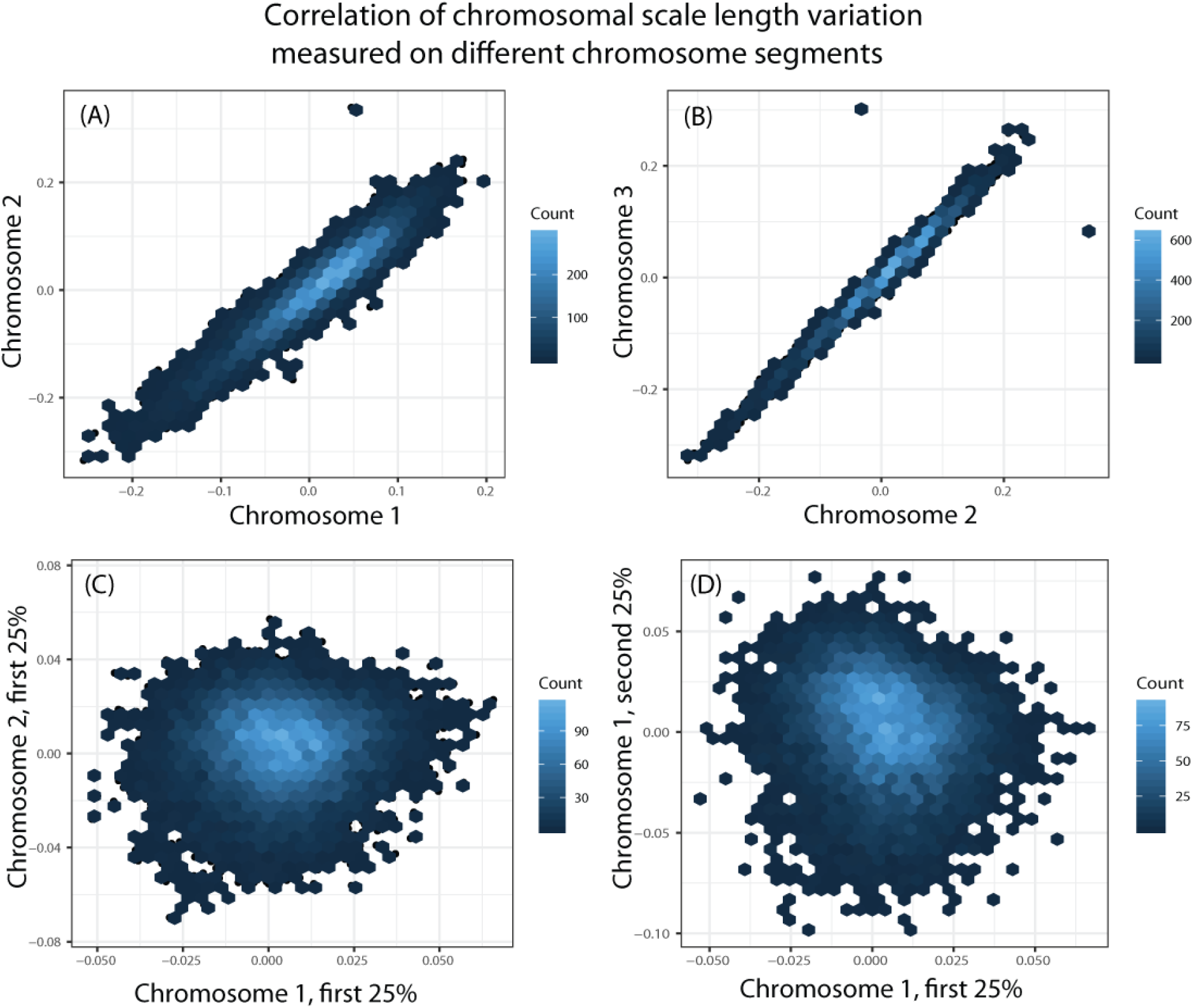
This figure depicts the correlation in chromosomal scale length variation measurements. We measured the chromosomal scale length variation for 10,000 different people in the UKBiobank dataset for different chromosomes and segments of chromosomes. We found that the length variation measurement was sometimes highly correlated between chromosomes, as in (A) and (B). However, when we measured the length variation in subsets of the chromosome, as in (C), we might find little correlation between the length variation measured on subsets of the chromosomes. In this figure, all axes have the units of the logarithm, base 2, of the average length change. A value of 0 indicates the person has a normal length of the chromosome segment. A value of 0.2 indicates the person’s chromosome is about 15% longer than normal.

## Discussion

Several different polygenic risk scores have been developed for different forms of cancer(17,18). Breast cancer was the first target of genetic risk scores with the discovery of BRCA1. It probably has the most advanced polygenic risk scores available today. The 2015 study that computed a polygenic risk score based on 77 SNPs found that women who scored in the top 1% had a three-fold increase in risk compared to a woman who scored in the middle quintile (6). For comparison, in the ovarian cancer prediction presented here 24 of 33 in the top 3% had ovarian cancer, while 2 of 33 in the middle 3% had ovarian cancer, for an approximate tenfold increase in risk. All 11 of the women scoring in the top 1% had developed ovarian cancer.

Machine learning techniques have been used to build polygenic risk scores to predict other complex traits(19–21). For instance, diabetes can be predicted from SNP data with an area under the receiver operator curve (AUC) of 0.602(21) using a gradient boosted regression tree.

We used the h2o platform for machine learning and selected the best algorithm by weighing computation time and AUC. The h2o platform tested four different algorithms (generalized linear model, distributed random forest, gradient boosting machine, and deeplearning). For the TCGA data, the gradient boosting machine algorithm and the deep learning algorithm provided comparable AUCs, but the gradient boosting algorithm was substantially faster. For the UKBiobank data, the deep learning algorithm provided substantially larger AUCs than all other algorithms tested.

The tests have not been optimized. The UKBiobank test could be further improved in two ways. First, the results could improve with further training. With a training time of ten hours, we obtained an AUC of 0.83 (range 0.82-0.84 with five replicates using different initial randomizations). Using only one-hour training, we could only obtain an AUC of 0.77 (range: 0.71-0.81 with 10 replicates). The training was done with a desktop computer (Intel i7-3770 with 4 cores, no GPU). We believe additional training time could improve the AUC. Second, we used 88 numbers to characterize each genome, splitting the 22 chromosomes into four equal parts. This was the only split we tried. We did not explore alternatives, because we did not have the computer power to do so.

Heritability measures how well one can predict a phenotype of an offspring based on the two parents(22). This definition is often referred to as “broad-sense heritability.” This heritability can be estimated from studies of monozygotic and dizygotic twins(1,23). A large gap exists between heritability measured from twin and family studies and the extent to which we can currently predict a phenotype from genetic data. This gap is called the missing heritability or phantom heritability(5,24). Current predictions are based on additive models of SNPs, this is known as “narrow sense heritability”. Models based on chromosomal-scale length variability should be better measures of broad sense heritability, since they include epistatic effects.

Twin studies assume that monozygotic twins are genetically identical. This assumption, however, does not hold for copy number variation(25,26). Substantial difference in copy number variation occurs after conception. Thus, predictions based on chromosomal-scale length variation measured in peripheral DNA could be even higher than estimated by heritability measured in twin studies.

Interpreting the predictions made here is difficult. Although risk prediction and association studies share common methods, the end goals differ. Association studies often try to identify alterations in specific genes that can be mechanistically tied to specific diseases. Risk prediction, however, is only concerned with maximizing the predictive power(18).

Some of our predictions are generated using gradient boosting machines, which are derived from decision trees(21,27). Decisions trees are easily interpretable. However, the useful variables are not. For instance, one point in the decision tree asks whether a certain measured chromosomal scale length variation, expressed as a segmented mean value of a 133 mega base region of chromosome 11 (specifically between bases 456,012 and 134,272,740 in hg38 coordinates) is greater than or less than 0.0090125. The segmented mean value of 0.0090125 represents an excess length of about 1% in that region. This is one of thousands of different decisions the algorithm examines to compute a prediction. Identifying the mechanisms behind these predictions is not simple, but something we plan on working towards.

Interpreting the area under the receiver operator curve (AUC), which is also known a C-index, is straightforward. Classification tests are a tradeoff between the true positive rate, or sensitivity, and the false positive rate, or 1-selectivity. These tests provide a numerical value. In practical use, a cutoff value must be selected, where all tests above the cutoff fall into one class and all below fall into another. The cutoff value is arbitrary; different cutoff values will give different sensitivity and selectivity values. The receiver operator curve summarizes the sensitivity and selectivity values for different potential cutoff values. The integral of this curve, known as the area under the curve (AUC) summarizes the performance of the test independent of the arbitrary cutoff value. An AUC value of 1.0 is perfection. The test classifies each case correctly. An AUC value of 0.5 is useless; it represents a random test.

Some example area under the receiver operator curve (AUC)s for diagnostic tests are: the BNP test for congestive heart failure AUC=0.91 (28), the PSA test for prostate cancer AUC=0.68 (29), and more recently the Cancer SEEK test, a “blood biopsy” to determine whether one of six types of cancer are present AUC=0.91 (30). Predictive tests generally have lower AUCs than diagnostic tests. A predictive test for coronary artery disease using genetic information has an AUC of 0.62, but when combined with conventional risk factors it can be increased to 0.68(31). For comparison, 7 of the 31 cancer tests shown here have an AUC greater than or equal to 0.8 and 15 of the 31 tests have an AUC greater than or equal to 0.75 suggesting that most of these tests will be at least as useful as the current predictive tests.

We considered whether the results were due to two common problems faced by GWAS studies: batch effects or population stratification. This analysis might be identifying batch effects rather than real effects (32), but it seems unlikely. First, all samples were collected from the same tissue, blood. This eliminates one common source of batch effects, since the DNA extraction process is the same for each sample. Second, all samples were processed by the same laboratory, the Nationwide Children’s Hospital Biospecimen Core Resource, with the same type of instrument. Most of the samples were processed in multiple batches (the number of batches is indicated in Table 2). Samples from five different types of tumor patients were processed in individual batches, but these were not outliers in the results. Replicating the results in an independent dataset, the UKBiobank dataset, further rules out batch effects.

Population stratification occurs in case/control studies when the cases and controls contain substantially different proportions of genetically discernable subclasses. Most TCGA samples were collected in the United States from a racially diverse group. For instance, over half the ovarian cancer samples were collected at five locations in the United States: Memorial Sloan Kettering, Washington University, University of Pittsburgh, Duke, and Mayo Clinic-Rochester. Table 3 outlines the breakdown. This table does indicate slightly different proportions, by race, in the case and control groups. The ultimate test of a predictive model is whether one derived from one group can applied to a different group. Table 4 demonstrates that the predictive model derived from a strictly white population works equally well, if not better, in an Asian population. It also performs good, but substantially worse in a black/African American population. This observation is consistent with our understanding of how genetic variation correlates with race(33,34).

Cancer is the result of a complex interaction between genetics and the environment. In some cases, for instance, lung cancer and smoking or mesothelioma and asbestos exposure, the required environmental exposure is significant and well known. In other cases, the required environmental exposure is probably minor and not well known. The genetic signature identified here is a necessary, but not sufficient factor in developing the cancer. Since this is a retrospective study of people who already developed cancer, sufficient environmental exposure has already occurred. A prospective study would need to be performed to determine the effect of environmental exposure on how effective these predictions are.

## Conclusions

In conclusion, we found that most cancers have significant inherited components. Machine-learning techniques applied to germline DNA chromosomal-scale length variation can effectively predict whether a person will develop many of these tumors. This method should be generally applicable to predict any cancers or other complex genetic conditions including conditions as diverse as heart disease and schizophrenia.

## Methods

The Cancer Genome Atlas (TCGA) was a project sponsored by the National Cancer Institute to characterize the molecular differences in 33 different human cancers (9,10). The project collected samples from about 11,000 different patients, all of whom were being treated for one of 33 different types of tumors. The samples collected usually included tissue samples of the tumor, tissue samples of normal tissue adjacent to the tumor and normal blood samples. (Normal blood samples were not available from patients diagnosed with acute myeloid leukemia.)

Most of the patient blood samples were processed to extract germline DNA. All TCGA germline DNA samples were processed by a single laboratory, the Biospecimen Core Resource at Nationwide Children’s Hospital. Single nucleotide polymorphisms (SNPs) were measured from the patient samples with an Affymetrix SNP 6.0 array. This SNP data was then processed (by the TCGA project) through a bioinformatics pipeline (35), which included the packages Birdsuite (36) and DNAcopy (37). The result of this pipeline is, for each sample, a listing of a chromosomal region (characterized by the chromosome number, a starting location, and an ending location) and the associated value given as the “segmented mean value.” The segmented mean value is defined as the logarithm, base 2 of one-half the copy number. A normal diploid region with two copies will have a segmented mean value of zero.

NCI has provided most of the TCGA data on the Genomic Data Commons (38). The copy number variation is called the masked copy number variation. The masking process removes “Y chromosome and probe sets that were previously indicated to have frequent germline copy-number variation.” (35)

The final data set that we used is the masked copy number variation data. It originates from normal blood samples extracted from 8,826 different patients: 4,692 female and 4,134 male. The patient’s age ranged from 10 to 90 years old.

In this dataset, about 695,000 different genomic regions characterized by the chromosome number, a start position, and an end position have additions or deletions, known as copy number variations, in at least one patient.

We initially selected the top 10,000 of these 695,000 regions, ranked by the number of different patient samples that contained that CNV, to create a dataset with 8,826 rows and 10,000 columns. After some work on this dataset, we noted that many overlapping CNVs exist. As an improved second version, we used an alternate grouping characterized simply by the chromosome number and the end location. This groups together all CNVs that share the same end location. This second version gave better results than the first version for most tumors. We used the second version throughout the manuscript.

For each type of tumor, we created a case/control study. Cases, shown in Table 2, were the number of “normal blood” samples from patients with those diagnoses. This quantity is the number of patients who also have measurements of chromosomal-scale length variation from DNA derived from normal blood in the database.

Controls, also shown in Table 2, were the number of “normal blood” samples from patients without that specific diagnosis. These normal blood samples included patients with other diagnosis. A few forms of cancer were specific to women (cervical squamous cell carcinoma, ovarian serous cystadenocarcinoma, uterine corpus endometrial carcinoma, and breast invasive carcinoma) so the controls only consisted of female controls. The male specific forms of cancer (prostate adenocarcinoma and testicular germ cell tumors) had only male controls. Nine cases of male breast cancer are included in the dataset, so breast cancer was analyzed both as female specific and for either sex. Each cancer test was set up as a binary classification supervised learning task that was trying to distinguish between patients diagnosed with one cancer (breast cancer, for example) and those not diagnosed with that cancer.

We performed the analysis using the statistical language R. The TCGA copy-number variation data was acquired through a query to Google’s BigQuery, which hosts a copy of the TCGA data as part of the Institute of Systems Biology Cancer Genomics Cloud(39). The data was reformatted for analysis, and then was fed into the machine-learning algorithm. The data included the patient’s sex, whether or not they had the particular cancer, and measurements of chromosomal-scale length variation at different locations derived from the patient’s peripheral blood sample. The data fed into the machine-learning algorithm did not include age, since germline DNA should not depend upon age. The results are independent of the patient’s age.

The machine-learning algorithm used the h2o platform, provided by the h2o library in R (http://www.h2o.ai). We initially used a deep learning network for the analysis and had good results without tuning any hyperparameters. We used the default values provided by h2o (two hidden layers, each with 200 neurons, the default learning rate, etc.). We tested several other algorithms. We generally obtained the best results using the gradient boosting machine algorithm, again without tuning any of the hyperparameters. The gradient boosting machine algorithm was considerably faster, so we present our results here with the gradient boosting machine algorithm.

We used 10-x cross-validation for the analysis of the TCGA data. The result of the machine learning analysis is a computational model that provides a “score,” indicating a likelihood of a patient having a cancer, given a list of chromosomal-scale length variations. The computational model can be characterized by the area under the curve of the receiver operator curve, commonly referred to as the AUC of the ROC. We recorded the AUC of the model and repeated the process five times. This provided a measure of the variance of the process.

An independent validation strategy gave similar results to the cross validation analysis. We split the data randomly into two sections, an 80% training set, and a 20% validation set. The training set was used to build the model without any input from the validation set. Once the model was built, it was applied to the validation set. The AUC for this strategy were similar to those observed in the cross validation study.

We also tested a random control. One third of the patients in the dataset were randomly labelled as cancer, while the other two thirds were labelled as healthy. Following the same independent validation procedure, we attempted to build a model that could predict to which category a patient should belong. For this random control, we found that the area under the receiver operator curve (AUC) was 0.50 with a standard deviation of 0.01. The classification was equivalent to random guessing.

### UKBiobank

We were granted access to UKBiobank data as part of application 47850.We downloaded the “l2r” data for all 22 chromosomes using the ukbgene tool provided by UKBiobank. The l2r data is the log, base 2, of the transformed intensity values measured for each of about 850,000 variants spread across the 22 chromosomes. These 850,000 measurements are recorded for each of the 488,377 participants in the study. The measurements are recorded in 22 separate files that, in total, are about 3 TB. The files are plain text, space-separated data. Each column represents a unique participant. Each row is a different SNP location on the genome.

We used the unix command line programs “cut” to extract relevant columns from the UKBiobank dataset and “split” to partition chromosomal data into four equal subparts. Once these large files were reduced to manageable size by “cut” and “split”, they were read into R and an average chromosomal scale length variability value was computed for each subject by taking the mean of all the l2r values in the file for the chromosomal segment. This process gave a set of 88 numbers (four for each chromosome segment multiplied by 22 different chromosomes) for each subject. These 88 numbers per subject were used for machine learning classification and for the correlation studies.

For the classification study, we selected 1,534 women who both self-reported having breast cancer and were recorded by a cancer registry as having breast cancer. These were all the cases fitting this criteria in the entire population of 488,377 people. We also selected 4,391 cancer free women as controls. These 4,391 were selected by starting with an arbitrary 10,000 people. We first removed all men from the set of 10,000 people. Then we removed any women from the set who had any record of any type of cancer. From these 4,391 controls and 1,534 cases, we constructed a labelled dataset of 5,925 subjects each characterized by 88 numbers. We used h20’s automl function to identify the best algorithm using 5x cross validation. We repeated the analysis five times with different random seeds to produce different cross-validation splitting.

## Data Availability

This paper is about analyzing previously collected data. The data we analyzed is available from TCGA at https://cancergenome.nih.gov/. The demo code will download a subset of the TCGA data necessary for the analysis in this paper. The UKBiobank data is not publicly. However, it is easily available following an application to UK Biobank.

## Code Availability

The code is written in R. It is available as a supplemental file to this paper.

## Acknowledgments

The results published here are in whole or part based upon data generated by the TCGA Research Network: http://cancergenome.nih.gov/. This research has been conducted using the UK Biobank Resource under application 47850..

